# Site-specific recombination between inverted target sites generates dicentric/acentric chromosomes

**DOI:** 10.1101/2020.05.27.112383

**Authors:** Simon W. A. Titen, Makenna T. B. Johnson, Mario Capecchi, Kent G. Golic

## Abstract

Site-specific recombinases are widely used tools for analysis of genetics, development and cell biology, and many schemes have been devised to alter gene expression by recombinase-mediated DNA rearrangements. Because the *FRT* and *lox* target sites for the commonly used FLP and Cre recombinases are asymmetrical, and must pair in the same direction to recombine, construct design must take into account orientation of the target sites. Both direct and inverted configurations have been used. However, the consequence of recombination between target sites on sister chromatids is frequently overlooked. This is especially consequential with inverted target sites, where exchange between oppositely oriented target sites on sisters will produce dicentric and acentric chromosomes. By using constructs that have inverted target sites in *Drosophila melanogaster* and in mice, we show here that dicentric chromosomes are produced in the presence of recombinase, and that the frequency of this event is quite high. The negative effects on cell viability and behavior can be significant, and should be considered when using such constructs.

## INTRODUCTION

Site-specific recombinases have become essential tools for biological experimentation. The FLP-*FRT* and the Cre-*lox* systems have been used to great effect in a number of animal and plant systems. Genes or gene segments may be arranged in a variety of configurations with *FRT* or *lox* target sites to activate or inactivate genes at different developmental stages in animal and plant systems. By using such methods, cells and their descendants may be marked to study the fates of mutant cells or to trace cell lineage. Additionally, recombination between widely separated target sites on the same chromosome, or between homologous or nonhomologous chromosomes, has been used to engineer a variety of chromosome rearrangements(Golic and Lindquist 1989; Golic 1991; Lakso *et al*. 1992; Qin *et al*. 1994; Smith *et al*. 1995; Ramírez-Solis *et al*. 1995; Golic and Golic 1996; Gilbertson 2003; Zong *et al*. 2005; Wang *et al*. 2010; Lee 2013; Gierut *et al*. 2014; Hubbard 2014; Weissman and Pan 2015; Pontes-Quero *et al*. 2017; Germani *et al*. 2018).

The *FRT* and *lox* target sites are both asymmetrical, and FLP- or Cre-mediated recombination generates orientation-specific outcomes. Intramolecular recombination between sites in the same (direct) orientation excises the material between the sites, along with one target site (**Figure 1A**). Intramolecular recombination between sites that are inverted results in an inversion of the material between them (**Figure 1B**). Intermolecular recombination may also occur, and the consequence of recombination between sister chromatids is often overlooked. Recombination between target sites at identical locations on sister chromatids is typically without effect (**Figure 1C**). Recombination between target sites at different locations, though, does produce genetic changes that may be significant. When two target sites are in the same orientation, recombination between staggered sites generates complementary duplications and deletions (**Figure 1D**). Recombination between inverted target sites on sister chromatids will generate a dicentric and an acentric chromosome (**Figure 1E**). If the inverted target sites are close to one another, this can be quite frequent. These outcomes are reversed if the target sites are on opposite sides of the centromere, but since the frequency of recombination is strongly dependent on proximity (Golic and Golic 1996), this is of much less concern.

**Figure 1.**
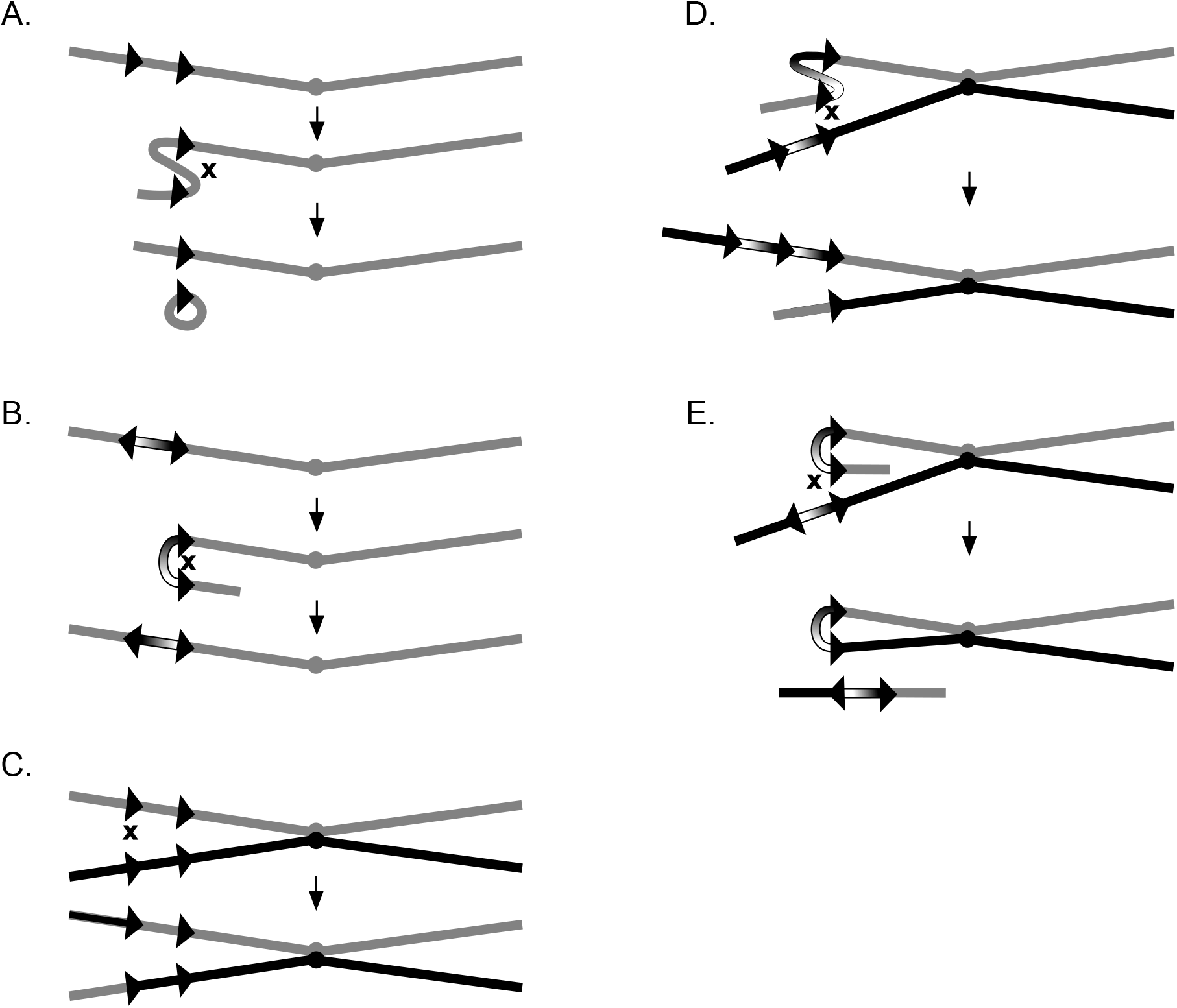
Intramolecular and intermolecular site-specific recombination. Schematic representations of recombination between recombinase targets on the same chromatid or on sister chromatids. (A) Intramolecular recombination between targets in direct orientation results in excision of the material between the targets as an extrachromosomal circle. (B) Intramolecular recombination between inverted repeats inverts the orientation of the DNA between the target sites. (C) Intermolecular recombination between targets at the same location on sister chromatids results in equal exchange. (D) Exchange between targets in direct orientation at different sites on sisters results in duplication and deletion of intervening DNA. (E) Recombination between inverted target sites on sisters produces a dicentric chromosome and an acentric chromosome. Chromatids, are indicated as lines, target sites as arrowheads and centromeres as filled circles.

In mitosis, a dicentric chromosome can break, leading to breakage-fusion-bridge cycles, chromosomal and segmental aneuploidy, and cell death. In cells that do not die, continuing problems with chromosome segregation and aneuploidy can cause aberrant behavior. Falco *et al*. (Falco *et al*. 1982) were the first to recognize the problem created by inversely oriented recombinase target sites within a chromosome. They saw that chromosomal integration of the 2μ plasmid of *Saccharomyces cerevisiae*, which carries inverted *FRTs* and encodes the FLP recombinase, caused chromosome instability, which they attributed to dicentric/acentric formation. Subsequently, this event was directly demonstrated in *Drosophila melanogaster* (Golic 1994), and inferentially shown in mice with Cre-*lox* (Lewandoski and Martin 1997). In the latter case, when inverted *lox* sites were placed on the mouse *Y* chromosome, and males with this *Y* were mated to females with a *ß-actin-Cre* gene, almost all *XY* zygotes developed as *XO* females, owing to loss of the *lox-*bearing chromosome during embryonic development.

Since those examples, there have been a number of studies that have utilized FLP- or Cre-mediated recombination between inverted target sites to cause dicentric chromosome formation (Ahmad and Golic 1998; 1999a; Otsuji *et al*. 2008; Grégoire and Kmita 2008; Titen and Golic 2008; Tada *et al*. 2009; Titen and Golic 2010; Jones *et al*. 2010a; Zhu *et al*. 2010; Kurzhals *et al*. 2011a; Zhu *et al*. 2012; Sato *et al*. 2017; Thomas *et al*. 2017). Other studies have identified cell death as a consequence of recombination between inverted target sites without directly pinpointing the cause (Eckrich *et al*. 2019).

In spite of these many examples of inverted target sites causing dicentric chromosome formation in the presence of the recombinase, their use for gene switching and lineage tracing continues, apparently without recognition that dicentric chromosomes are formed. Flex and FlipFlox constructs, used for gene switching in mice, rely upon Cre-mediated inversion (Schnütgen *et al*. 2003; Xin 2005). The poly*lox* system for barcoding cells and tracing cell lineage in mice relies on both direct and inverted *lox* repeats (Pei *et al*. 2017). The Confetti method for cell marking, and other methods based on Brainbow 2.0 or 2.1, also utilize inverted target sites (Livet *et al*. 2007; Snippert *et al*. 2010; Richier and Salecker 2014; de Roo *et al*. 2019). In flies, Flybow, FlpStop, Flip-Flop and other implementations have used inverted target sites to alter gene expression during development (Hadjieconomou *et al*. 2011; Chin *et al*. 2014; Fisher *et al*. 2017; Nagarkar-Jaiswal *et al*. 2017; Williams *et al*. 2019). In yeast, the SCRAMBLE system for rearranging chromosomes uses engineered symmetrical *lox* sites that may recombine in either direction (Dymond *et al*. 2011; Annaluru *et al*. 2014; Wu *et al*. 2018).

Studies that use these constructs will be impacted by the death of cells with broken chromosomes, or by the persistence of cells with aneuploidy and continuing genome instability, possibly leading to misinterpretation or invalid conclusions. Our purpose here is to raise awareness of the fact that dicentric chromosomes are formed when inverted recombinase target sites are used in eukaryotes, and that this event can be very frequent. This outcome should be considered when planning experiments with such constructs.

## RESULTS

As a practical demonstration of dicentric chromosome formation, we obtained flies carrying an insertion of the Flip-Flop construct, either on chromosome *2* or chromosome *3*. The Flip-Flop cassette was developed as a method for switching between GFP and mCherry labelling of gene traps in Drosophila (Nagarkar-Jaiswal *et al*. 2017).

Inverted *FRTs* were placed so that inversion of the flanked segment switches between the two markers. To test whether dicentric chromosomes would form after expression of the FLP recombinase, we generated progeny that contained the Flip-Flop construct and a heat shock inducible FLP gene (*70FLP*). We heat shocked larvae for one hour at 38° and then examined neuroblast mitotic spreads from third instar larvae (**Figure 2**). With a Flip-Flop insertion on *2L*, 63% of metaphases showed dicentric/acentric chromosome *2* figures at 4-5 hrs after heat shock. On *3R*, 46% of chromosome *3* figures showed the same. At 23 hrs after heat shock, cells showing dicentric/acentric figures were approximately half as frequent as the earlier time point, owing, no doubt, to the arrest and death of cells suffering from aneuploidy and broken chromosomes during the intervening period (**Table 1**; (Titen and Golic 2008)). These results demonstrate that FLP recombinase does not benignly switch markers within the Flip-Flop cassette, but also causes dicentric chromosome formation. Our previous work shows that this is frequently, but not inevitably, followed by apoptosis (Golic 1994; Titen and Golic 2008; Kurzhals *et al*. 2011a).

**Figure 2.**
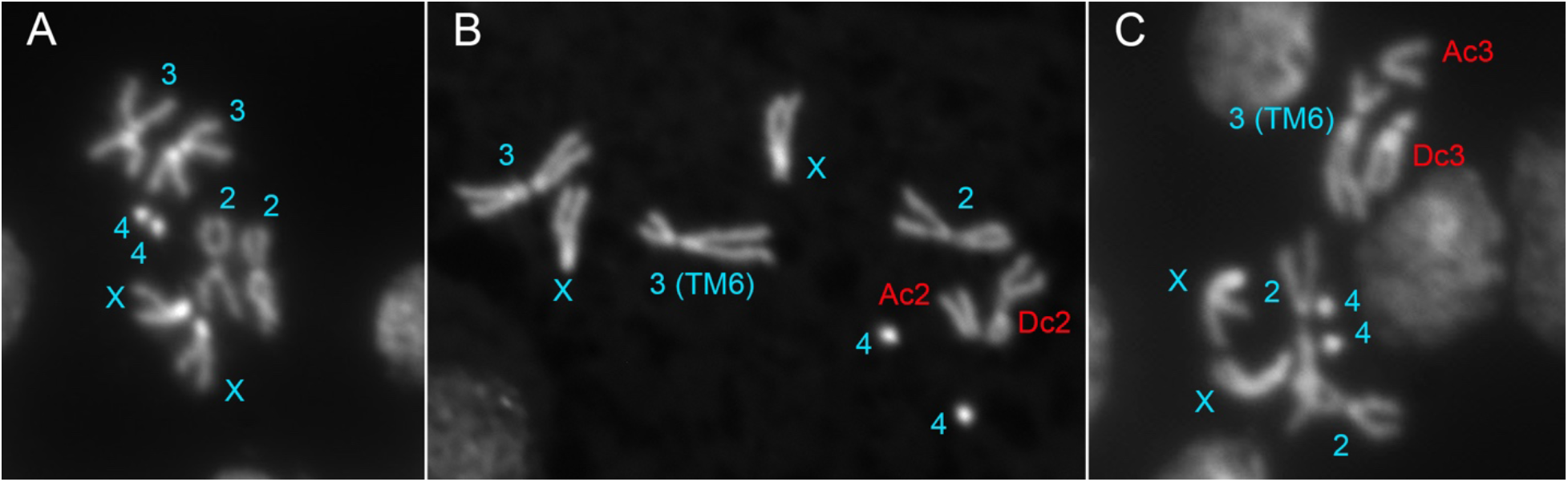
Dicentric chromosomes in Drosophila larval neuroblasts. (A) The normal *D. melanogaster* karyotype. (B) Dicentric and acentric chromosomes formed by unequal sister chromatid exchange using a FlipFlop construct inserted at 37B on chromosome *2*. Normal chromosomes are indicated in blue; dicentric (Dc) and acentric (Ac) products indicated in red. (C) Dicentric/acentric chromosomes produced using a FlipFlop construct at 82E on chromosome *3*.

**Table 1.**
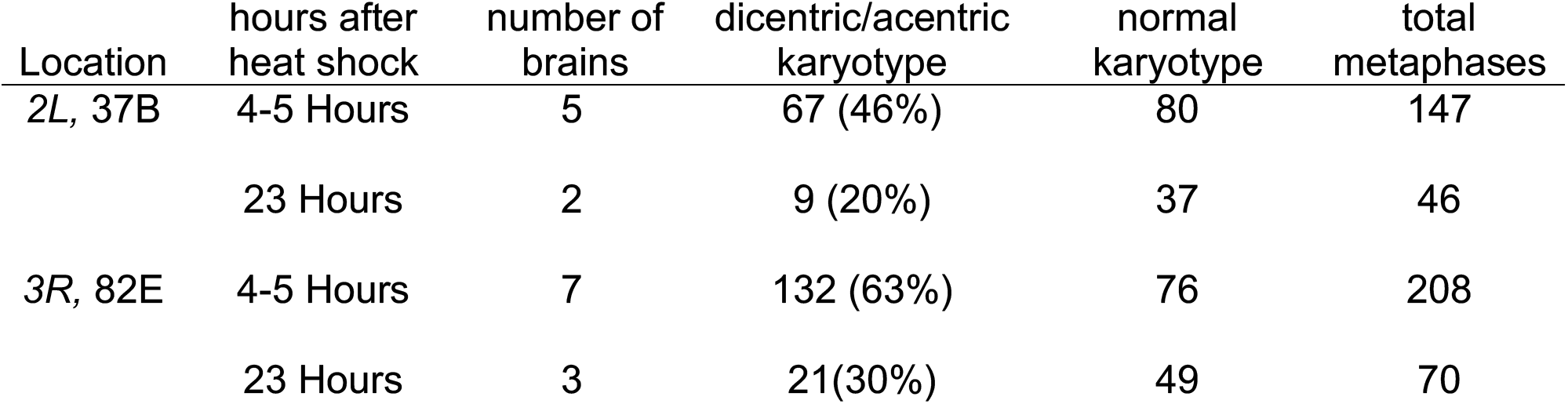
Dicentric chromosome frequencies with FlipFlop insertions.

In the mouse, there is also ample evidence that the use of inverted *lox* sites produces dicentric chromosomes. As a direct demonstration of this, we generated Embyonic Stem (ES) cells carrying a targeted insertion of a construct with inverted *loxP* sites telomere adjacent on chromosome *7* within the coding region of *fgf4*, and Mouse Embryonic Fibroblasts (MEFs) with inverted *loxP* sites near the centromere of chromosome *15* within the coding region of *ext1*. The ES cells were transfected with Cre recombinase expression vector and the MEFs were treated with the Cre^TAT^ protein on plates. In either case we would expect to see splitting of the targeted chromosome into acentric and dicentric portions (**Figure 3**). Following incubation in the presence of the Cre recombinase we observed dicentric and acentric chromosome formation (**Figure 4A-D**; **Table 2**). Upon extended culture we observed further chromosome rearrangements, polyploidy, and evidence of breakage-fusion-bridge cycles (**Figure 4E-H**). We also directly visualized dicentric chromosome formation in live embryos carrying the inverted *loxP* sites on the tip of chromosome *7* and an *HPRT^CRE^* transgene — a constitutive *Cre* allele whose expression results in maternal deposition of the Cre protein (**Figure 5**).

**Figure 3.**
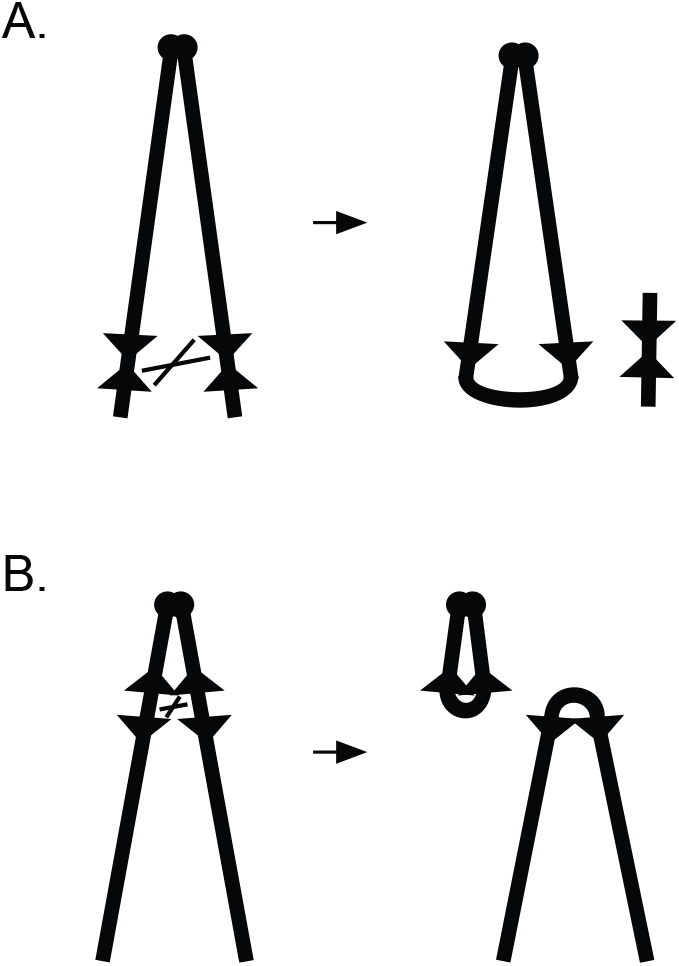
Dicentric chromosomes produced in mouse cells. Following exposure to Cre, asymmetric recombination between inverted *loxP* sites targeted to a locus near (A) the end of chromosome *7*, or (B) near the centromere of chromosome *15*, would result in a dicentric chromosome and an acentric chromosome fragment. As indicated, the recombinant chromosome products are expected to be significantly different in size.

**Figure 4.**
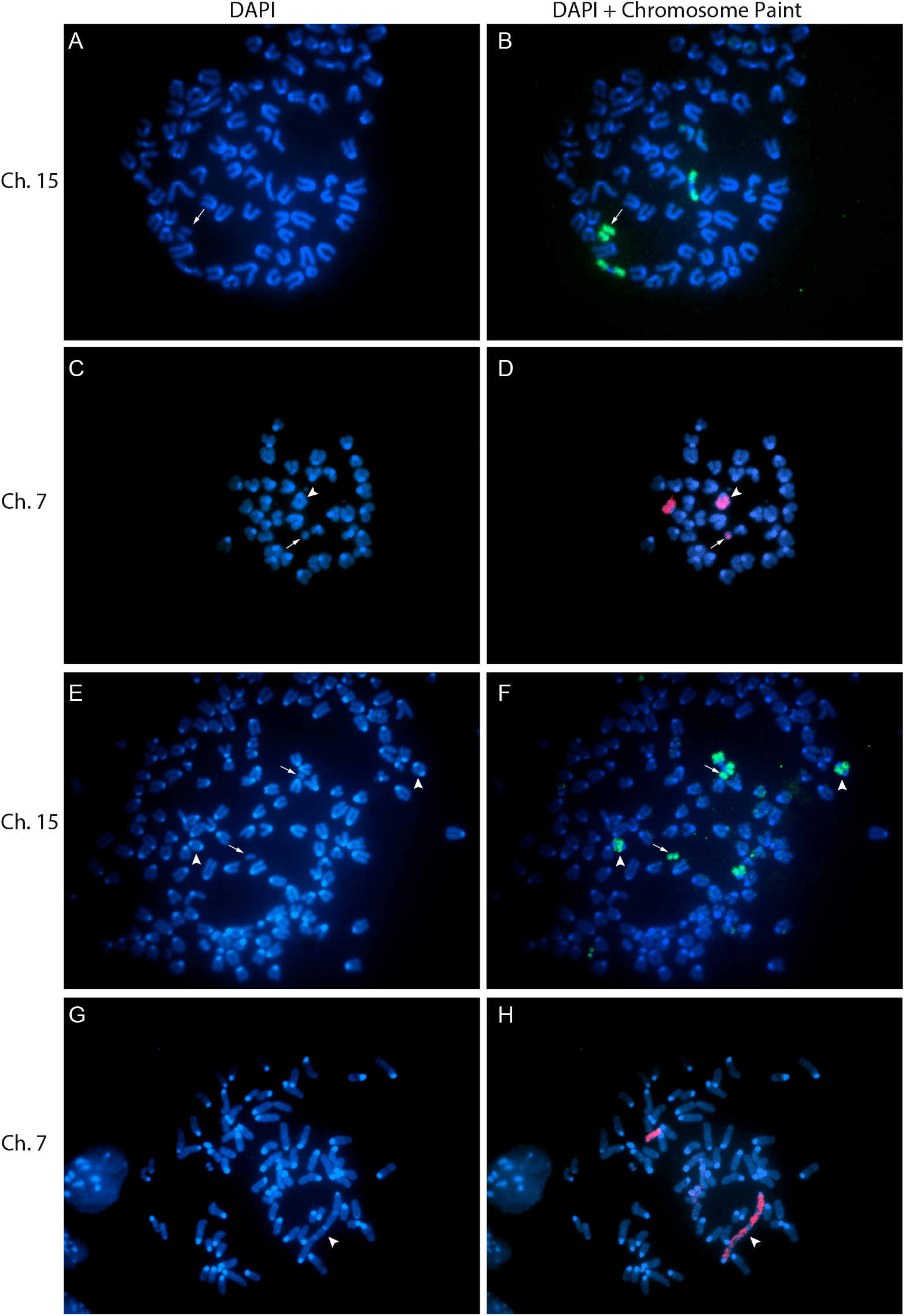
Evidence of Cre-mediated dicentric and acentric chromosome formation in mouse cells. Pictured here are mitotic figures of cultured cells transfected with a Cre expression vector or purified Cre protein. The targeted chromosomes were visualized with DAPI (A, C, E, G), or DAPI plus whole chromosome paints (B, D, F, H) specific for either chromosome *15* (A, B, E, F) or *7* (C, D, G, H). Exposure to Cre for a short duration (~24 hrs) results in large acentric fragments (A and B) or short acentric fragments and a large dicentric chromosome (C and D). Longer exposure to Cre results in evidence of further chromosome damage, such as smaller than expected chromosomes (E, F) or extraordinarily long chromosomes (G, H). Arrows indicate acentric fragments and arrowheads indicate dicentric chromosomes.

**Figure 5.**
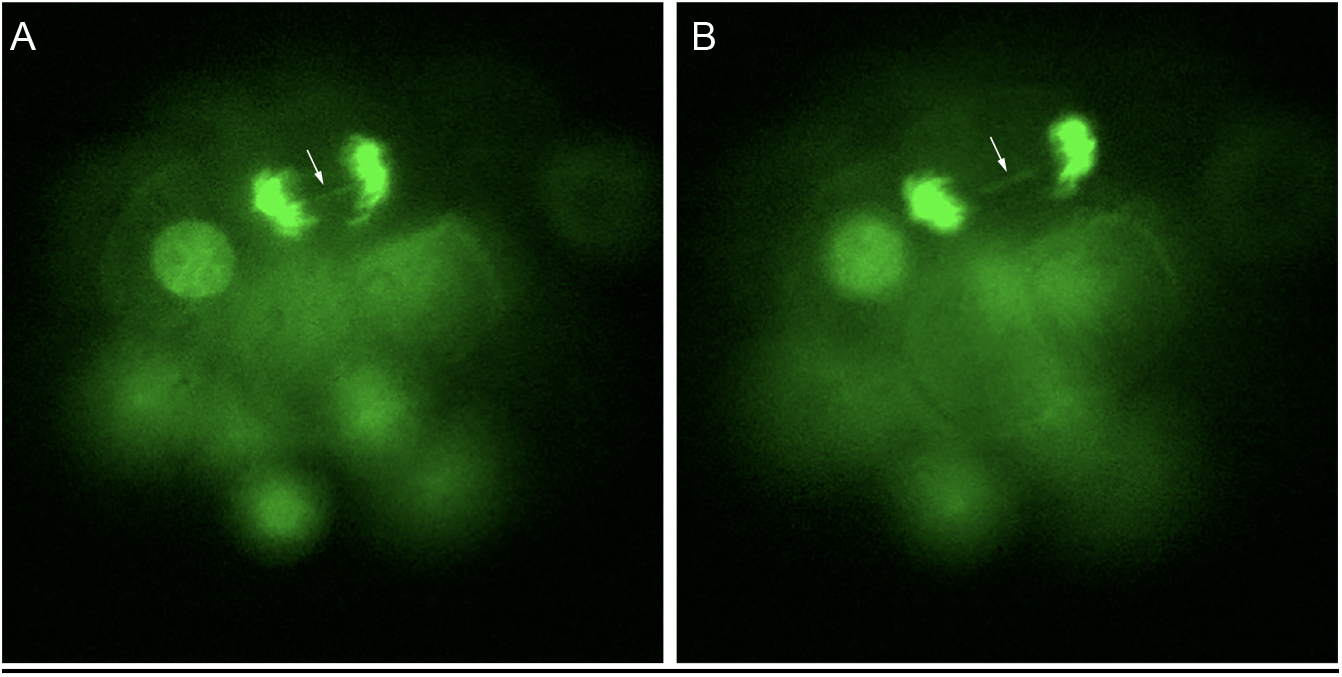
Dicentric bridge formation in early embryos. Mouse embryos carrying two inverted *loxP* sites on chromosome *7* with maternal Cre and Histone-GFP were used to make time-lapse movies of the early cleavage divisions. A and B are images of two successive timepoints during anaphase showing persistence of a dicentric chromosome during mitotic anaphase (arrow).

**Table 2:** Dicentric chromosome production in mouse cells. Cells with inverted *loxP* targets, inserted on chromosome *7* or chromosome *15*, were exposed to Cre protein and scored for dicentric or acentric figures, or polyploidy (>2N), in metaphase. Chromosome-specific paints were used to recognize chromosome *7* or *15*.

| insertion<br>site | normal<br>karyotype | dicentric<br>chromosome | acentric<br>chromosome | >2N | n optical<br>fields |
| --- | --- | --- | --- | --- | --- |
| chr. 7 | 40 | 15 | 73 | 16 | 100 |
| chr. 15 | 40 | 19 | 23 | 19 | 29 |

As a second test of dicentric/acentric formation in live embryos, we generated mice carrying a construct with the *Tet Repressor* (*Tet^R^*) gene distal to inverted *loxP* sites inserted in the subtelomeric region of chromosome *19*. Proximal to the *loxP* sites is a tetracycline-repressible reporter gene encoding the red fluorescent protein tdTomato (**Figure 6**). When dicentric/acentric chromosomes are formed by Cre activity, and the acentric portion carrying the *Tet^R^* is lost, *tdTomato* can be expressed. We examined 2-32 cell early mouse embryos carrying this construct and *HPRT^Cre^*. We observed *tdTomato* activation in a subset of cells, indicating that *Tet^R^* had been lost (Figure 6).

**Figure 6.**
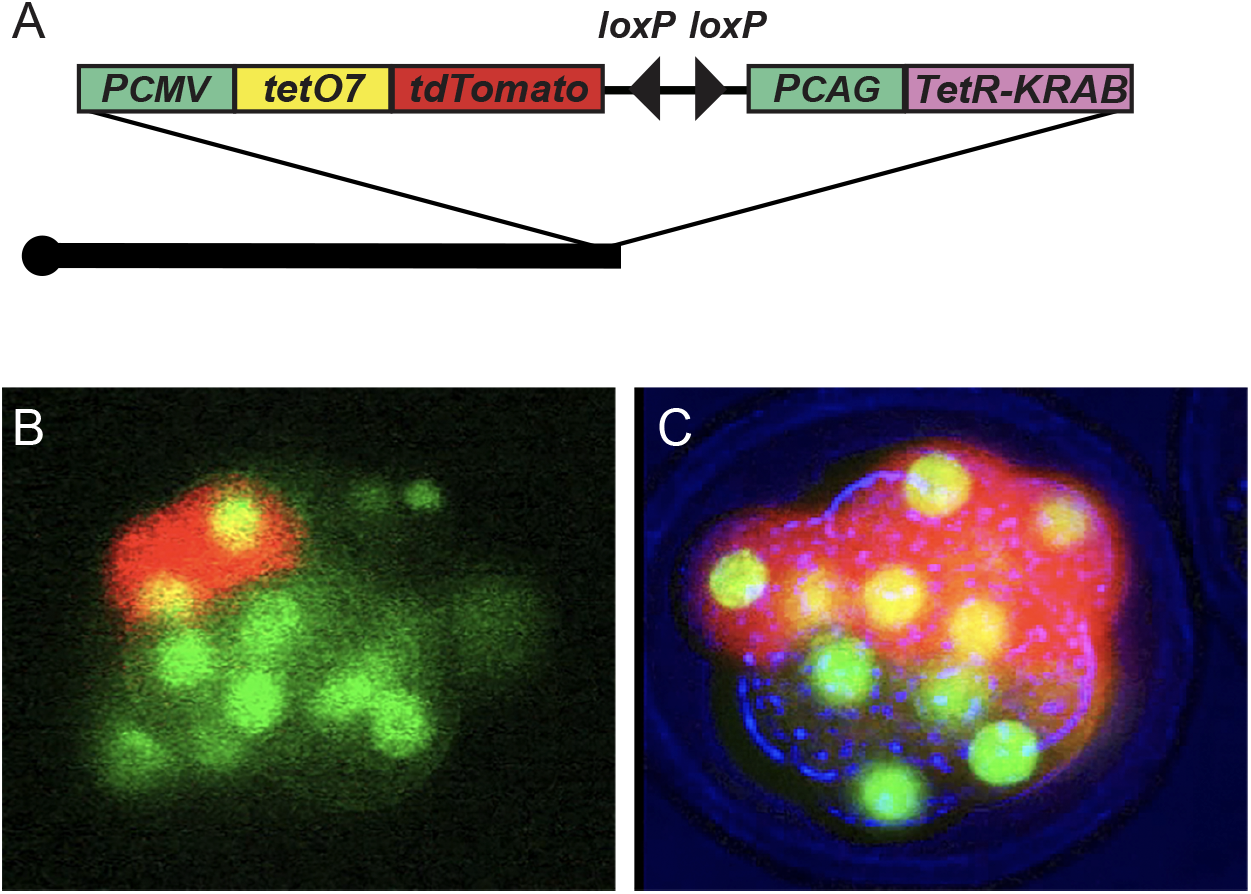
Marking cells after loss of acentric chromosome. (A) Schematic representation of the transgene targeted to the tip of *chromosome 19*. The transgene encodes a minimal *CMV* promoter (*Pcmv*), the tetracycline operator sequence (*tetO7*), and the *tdTomato* coding sequence, centromere proximal to inverted *loxP* sites; a strong *CAG* promoter (*Pcag*) controlling expression of the *Tet^R^-KRAB repressor* gene lies distal to the *loxP* sites. Loss of the *Pcag-tet^R^-KRAB* portion relieves repression of *tdTomato* expression, resulting in cells that fluoresce red. Early embryos carrying this construct, and maternally deposited Cre protein, were examined for tdTomato fluorescence. Nuclei are visualized with Histone-GFP (green). Cells that have lost the *Tet^R^* gene express tdTomato (red), two cells in (B) and ~6 cells in (C).

## Discussion

The occurrence of dicentric/acentric chromosomes will almost certainly confound the interpretation of experiments that use inverted target sites. We have shown, here and previously, that generation of dicentric/acentric chromosomes can be very frequent — in some cases nearly 100% of cells that express FLP and carry inverted *FRTs* can experience this event (Titen and Golic 2010; Kurzhals *et al*. 2017). These cells frequently, but not inevitably, undergo apoptosis (Ahmad and Golic 1999b; Titen and Golic 2008; Kurzhals *et al*. 2011a). This outcome can also be quite frequent when using Cre-*lox* in mice (Grégoire and Kmita 2008).

Experimental outcomes may be impacted, and possibly misinterpreted, by dicentric/acentric chromosome formation. As one possible example, recombination with the Confetti construct was used to mark and examine cell clones in the mouse intestine. Death of some clones, and their replacement by other clones, was observed (Snippert *et al*. 2010). It is difficult to exclude the possibility that this outcome was influenced by dicentric/acentric formation and subsequent death of cells with broken chromosomes or aneuploidy. More recently, a Brainbow construct was injected into single cell zebrafish embryos, along with a Cre expression vector, and the multicolor output was used to trace cell fate in the developing brain. The authors observed death of clones of cells during brain development (Brockway *et al*. 2019). Although the particular Brainbow construct used carries only directly repeated *lox* sites, the method of transformation typically generates tandem arrays of the injected DNA, and in many, if not most cases, some of the inserted copies are in opposite orientation to the others (Stuart *et al*. 1988; 1990; Culp *et al*. 1991). In this case, the formation of dicentric/acentric chromosomes is likely. Although it is possible that clonal death is a normal part of development of the central nervous system, in this experiment it may instead be a consequence of the formation of dicentric and acentric chromosomes. In this context, it is worth noting that cells which experience dicentric/acentric formation can divide several times to produce a clone before they die. Such cells can even differentiate into adult tissues, though they are less successful than wildtype cells (Golic 1994; Otsuji *et al*. 2008; Titen and Golic 2008; Tada *et al*. 2009; Zhu *et al*. 2010; Kurzhals *et al*. 2011b).

Our results, and those from other groups, make it exceedingly clear that the use of inverted target sites for a site-specific recombinase will frequently lead to the production of dicentric and acentric chromosomes, with significant effects on the behaviors and fates of cells in which they occur. There are schemes that use directly repeated target sites for gene switches and for lineage tracing (Nagy 2000; Kwan 2002; Zong *et al*. 2005; Gierut *et al*. 2014; Hubbard 2014; Richier and Salecker 2014; Weissman and Pan 2015; Muñoz-Jiménez *et al*. 2017; Pontes-Quero *et al*. 2017; Germani *et al*. 2018). These do not suffer the drawback of generating dicentric and acentric chromosomes and can be applied to many experimental situations.

## Material and Methods

Dicentric chromosomes in *Drosophila melanogaster:* Flies were maintained on standard medium at 25°C. The *70FLP* gene has been previously described(Golic *et al*. 1997). FlipFlop stocks were obtained from the Bloomington, IN, USA, Drosophila Stock Center. Female *y w 70 FLP3F; Sb/TM6,Ubx* flies were crossed to male *y w; Mi{FlipFlop} Nedd8 [MI13776-FF.PT-EGFP] /CyO* (Bloomington 76596) or male *y w; Mi{FlipFlop} Cdep[MI12769-FF.PT-EGFP]* (Bloomington 76595). Parental flies were removed from the vials after seven days and vials were placed into a circulating water bath for one hour at 38° C to induce FLP. Larval brains were dissected either 4-5 hours after heat shock, or 23 hours after heat shock. Neuroblast chromosomes were prepared as described (Fanti and Pimpinelli 2004) and then stained with DAPI. Multiple brains were scored, and each clear mitotic figure was scored as showing acentric/dicentric chromosomes or unaffected normal chromosomes. Nuclei in which all chromosomes were not visible or could not be distinguished were not scored.

Generation of Inverted *loxP* transgenic mice: The *HPRTC^re^* constitutive and germline Cre expression lines were obtained from Jackson Laboratories. The mouse line bearing inverted *loxP* sites targeted to the middle of chromosome *15*, at the *Ext1* locus, is described in (Jones *et al*. 2010b). The mouse line bearing inverted *loxP* sites targeted to the distal tip of chromosome *7*, to the *Fgf4* locus, is described in(Moon *et al*. 2000), except that the *loxP* site in the 5’ UTR is inverted with respect the other *loxP* site. The mouse line with inverted *loxP* sites and the tetracycline-repressible *tdTomato* transgene targeted to the tip of chromosome *19* was generated using homology arms with sequence similarity to the region distal to the last exon of the coding region of *Grk5*. Briefly, this construct contains a minimal *CMV* promoter immediately upstream of 7 copies of the *tetracycline operon operator* sequence immediately upstream of the coding sequence of a tandem dimer of the *tomato fluorescent protein* and a polyadenylation sequence. Downstream of tomato are the two inverted *loxP* sites followed by a *CAG* promoter driving expression of the fusion protein consisting of *Tet^R^* fused to the strong transcriptional silencing *Kruppel-associated box domain* (*Tet-KRAB*).

Mouse metaphase Chromosome Analysis: For preparation of mitotic figures MEF and ES cell lines cells were grown to 50-70% confluence in DMEM supplemented with Na Pyruvate, NEAA and 10% FBS for MEFs, or DMEM supplemented with Na Pyruvate and NEAA and 15% FBS and LIF for ES cells. MEFs were synchronized by double block with 50ng/ml methotrexate and 1mM thymidine, and then arrested in metaphase with 1ug/ml colcemid. ES cells were arrested at metaphase with colcemid at 100ng/ml (Detailed protocol available upon request.)

Cre^TAT^ treatment: Rapidly dividing cells were grown to 50-70% confluence. Cells were then washed with PBS and incubated in serum-free media supplemented with 4uM Cre^TAT^ protein for 1 hour at 37°C.

## Acknowledgments

This work was supported by grants R01 GM065604 (KGG) and R01 MH093595 (MC) from the National Institutes of Health and a grant from the Halt Cancer at X Foundation (SWAT).

